# Illuminating the Druggable Genome through Patent Bioactivity Data

**DOI:** 10.1101/2022.07.15.500187

**Authors:** María Paula Magariños, Anna Gaulton, Eloy Félix, Tevfik Kizilören, Ricardo Arcila, Tudor Oprea, Andrew R. Leach

## Abstract

The patent literature is a potentially valuable source of bioactivity data. The SureChEMBL database (https://www.surechembl.org/) is a publicly available large-scale resource that contains compounds extracted on a daily basis from the full text, images and attachments of patent documents, through an automated text and image-mining pipeline. In this paper we describe a process to prioritise 3.7 million life science relevant patents obtained from SureChEMBL, according to how likely they were to contain bioactivity data for potent small molecules on less-studied targets, according to the classification developed by the Illuminating the Druggable Genome (IDG) project. The overall goal was to select a smaller number of patents that could be manually curated and incorporated into the ChEMBL database. We describe the approach taken, the results obtained, and provide some illustrative examples.

## Introduction

One of the most useful and compelling pieces of evidence for the druggability of a new biological target is the existence of molecules that bind with sufficient affinity to modulate the biological activity of the target. However, only about 11% of the proteome has either an approved drug or a compound known to modulate it^1^. Chemical probes represent a special type of small molecule for use in target validation studies, not only having good activity against the target but also selectivity, cellular activity and potentially other relevant criteria^2,3^ and are subjected to a peer review process to ensure the quality of any conclusions when used by the wider community.

The availability of open-access, public databases such as ChEMBL^4^ has greatly simplified the task of identifying potential molecules, by providing easy access to data curated from the peer-reviewed scientific literature. ChEMBL currently contains more than 18 million bioactivity data on almost 2 million compounds. The data in ChEMBL is extracted manually by curators from several core Medicinal Chemistry journals or deposited by experimental groups. Bioactivity data are also periodically exchanged with other databases, including PubChem BioAssay^5^ and BindingDB^6^.

An additional and potentially valuable source of information and data on bioactive molecules is the patent literature. In drug discovery, patents are routinely filed to protect novel inventions, by both industrial organisations and academic institutions. The relationship between the patent and the “traditional” (academic journal, peer-reviewed) literature has been examined in a number of published studies, mostly focussed on key questions relating to overlap/duplication and publication date. For example, in a 2009 paper the authors found that just 6% of compounds in patents also appeared in the scientific literature in one of the commercial sources included in their study (GVKBIO)^7^. A later study reported that only 10% of a set of compound-target pairs could be found in literature publications within one year of being reported in patents, and that on average data appears 3 to 4 years earlier in patents than in the scientific literature^8^. For compounds that progress into clinical trials and are ultimately approved, the gap between publication of the patent and publication of the compound in the medicinal chemistry literature can be even longer^6^. Another study concluded that the first molecules for a novel target tend to be published first in the literature, whereas novel small molecules (regardless of targets) more frequently appear first in patents than in literature^9^.

These and similar studies suggest that the patent corpus potentially represents a wealth of information that is not available elsewhere and/or may appear in the scientific literature only after a significant time delay. Previous studies that have attempted to search or annotate pharmaceutical patents with target-compound information include Akhondi *et al*. who produced a gold standard corpus of 200 patents annotated with chemicals, diseases, targets and modes of action, with the goal of benchmarking patent text mining performance^10^; Suriyawongkul *et al*. who focused on identifying targets in patent titles that contained bioactive compound information^11^; Tyrchan *et al*. who compared different methods to extract key compounds using structural information, and then applied one of the methods to the design of AXL kinase inhibitors^12^; Fechete *et al*. who searched full-text patents using keywords related to diabetic nephropathy and further narrowed the search by rules related to frequency and/or patent section of the keywords found, and subsequently extracted the genes mentioned in the claims section of these patents^13^; and Gigani *et al*. who performed a search in the SureChEMBL database^14^ using keyword and/or chemical structure searches, with the goal of identifying patents with compounds that could activate the BK_Ca_ channel^15^. These, and similar studies, also confirm that extracting information from patents poses many challenges, related to the length of the documents, writing style, structure, and content, among others^16,17^.

Patent data are currently available from a number of resources, including Google Patents (https://patents.google.com/), Lens (https://www.lens.org/), Espacenet (https://www.epo.org/searching-for-patents/technical/espacenet.html) and FreePatentsOnline.com. All of these systems allow searching for patents using various criteria. Pubchem provides links to WIPO PATENTSCOPE, a patent database, for every compound contributed by WIPO to PubChem (16 million compounds). This allows users to search for patents associated with each chemical structure. BindingDB includes a curated set of US granted patents, from which protein-compound activity data is extracted.

In the work reported here, we use SureChEMBL^10^ (https://www.surechembl.org/), which is a fully automated, chemical-structure-enabled database providing the research community with open and free access to the patent literature. Currently, SureChEMBL sources data from both patent applications and granted patents via full text patents from the US, European and World patent offices and titles and abstracts from Japan. SureChEMBL currently contains ∼140 million patents with ∼50,000 added monthly. Of these, ∼50 million patents are chemically annotated. Approximately 80,000 new compounds are extracted and added each month to the SureChEMBL chemistry database which currently contains more than 20 million unique structures. The data in SureChEMBL can currently be accessed via a web interface that enables users to perform text and chemical structure queries, filter the output and then display the results. The complete set of chemical structures and patent associations are also available for download.

The US National Institutes of Health established the Illuminating the Druggable Genome (henceforth IDG) project in 2014 (https://commonfund.nih.gov/idg), with the goal of improving our understanding of understudied proteins that belong to well-studied protein families, such as ion channels, GPCRs and protein kinases. One of the key deliverables from the IDG project is an informatics platform, Pharos (https://pharos.nih.gov/) that provides researchers with free access to relevant data on targets. An important aspect of the IDG project (and the data in Pharos) is the classification of human proteins into four Target Development Level (TDL) families, based on how well studied these proteins are^18^. In the Tclin category are targets of at least one approved drug; Tchem targets do not have approved drugs but they are modulated by at least one small molecule with a potency above the cut-off specified for the target protein family (≤30 nM for kinases, ≤100 nM for GPCRs and nuclear receptors, ≤10 μM for ion channels and ≤1 μM for other target families); Tbio do not have chemistry qualifying for the Tclin/Tchem categories but have a reasonable amount of information (e.g., a confirmed Mendelian disease phenotype in the Online Mendelian Inheritance in Man (OMIM) database^19^, publications, GO annotation, antibody reagents); while Tdark are understudied proteins with little annotation^16^. The full description of these categories can be found at http://juniper.health.unm.edu/tcrd/. Of particular relevance to the work here is the availability of small molecule modulators for new targets, consistent with other work suggesting that the lack of high-quality chemical probes for understudied targets is an important cause for lack of interest ^16,20^. The default IDG process uses ChEMBL^4^, Guide to Pharmacology^21^ and DrugCentral^22^ to identify bioactive compounds for a target and so to determine whether it can be included in the Tchem category. At the time of writing, Tclin proteins represent ∼3% of the human proteome; Tchem proteins represent ∼8%; Tbio proteins represent ∼58%; and Tdark proteins represent ∼31% ^3^.

In this paper, we describe methods to systematically mine the SureChEMBL^10^ patent corpus to identify new bioactivity data for Tdark/Tbio targets, with the aim of 1) including the bioactivity data in the ChEMBL database and 2) promoting some of these targets to IDG Tchem status. We quantify the added value of such methods in terms of the numbers of targets that are so “promoted” and then provide some specific illustrative examples.

## Methods

The starting point for our work was the set of patents extracted from SureChEMBL covering the years 2012 to 2018, flagged as life-science relevant according to the IPC codes^23^ or CPC codes^24^ present in the patents, for a total of 3.7 million patents (the specific codes used were A01, A23, A24, A61, A62B, C05, C06, C07, C08, C09, C10, C11, C12, C13, C14, G01N). These codes classify the patents into different areas of technology. The goal was to find patents with bioactivity data of small molecules against understudied targets (Tdark or Tbio categories according to the IDG classification).

Firstly, in order to determine which patents were likely to have bioactivity data, the files corresponding to the patents were processed to identify tables containing the following keywords:

IC50, XC50, EC50, AC50, Ki, Kd, pIC50, pXC50, pEC50, pAC50, -log(IC50), -log(XC50), -log(EC50), -log(AC50), concentration to inhibit, IC-50, XC-50, EC-50, AC-50, IC 50, XC 50, EC 50, AC 50.

69,289 patents (2%) out of the 3.7 million were thus flagged as potentially containing bioactivity data in tables.

Separately, we identified patents that might contain information about IDG Tbio and Tdark targets. A list of Tdark and Tbio IDG target names and gene symbols was obtained from the Target Central Resource Database (36,044 target names/symbols). We searched for these target names and their gene symbols in the patent titles, abstracts, descriptions and claims sections, in the context of specific phrases that could indicate bioactivity data of small molecules against them, for example: “inhibitor/s of x”, “activity of x”, among others. The combination of these 2 procedures allowed us to classify the patents into 6 groups: patents with tables and bioactivity keywords, and targets mentioned in titles or abstracts; patents with tables and bioactivity keywords, and targets mentioned in descriptions or claims sections; patents without tables with bioactivity keywords, and targets mentioned in titles or abstracts; patents without tables with bioactivity keywords, and targets mentioned in descriptions and claims; patents with tables and bioactivity keywords but no targets; and patents without tables with bioactivity keywords and without targets (figure 1).

**Figure 1.**
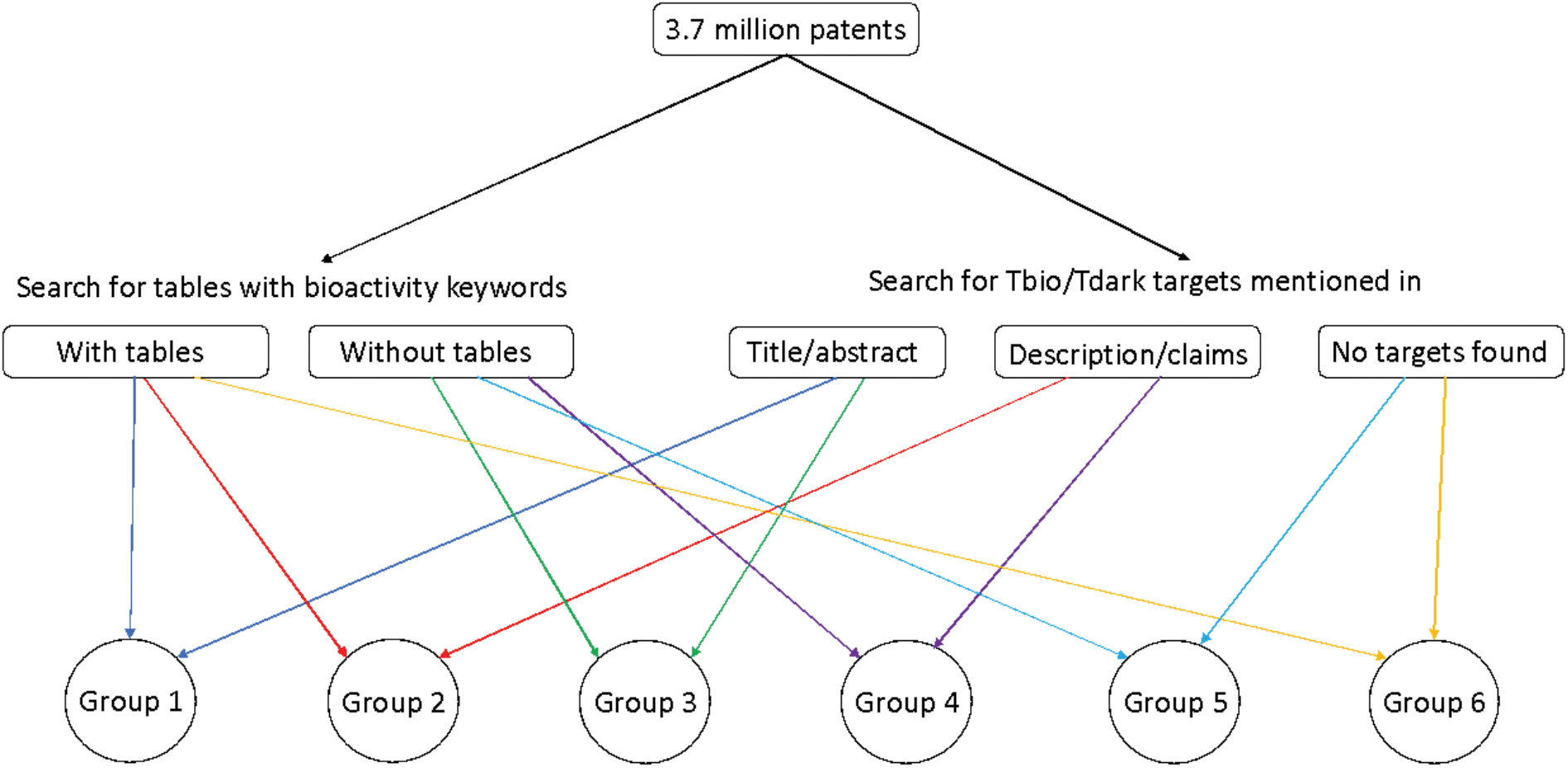
Overview of the classification of patents into six different groups, prior to further processing.

Following this automated annotation/filtering process, patents from each group were manually examined to confirm the presence of the correct Tbio/Tdark target, the presence of quantitative bioactivity measurements, and that the Tbio/Tdark target was the molecular target to which these bioactivity measurements applied. This final check is required because some of the patents were found to have data only on targets that did not belong to the IDG list of understudied targets; other patents did contain data exclusively on the targets of particular interest to us. Some patents fell into both categories.

For patents with confirmed bioactivity data, details of compounds synthesised, biological assays performed, and bioactivity measurements were manually extracted, according to the standard ChEMBL curation procedure described previously^25^ and loaded into the ChEMBL database. All bioactivities against all the targets present in these patents (irrespective of their inclusion or not in the IDG Tbio/Tdark categories) were extracted by the curators.

## Results

As a result of this work, bioactivity data from 225 patents were loaded into ChEMBL, corresponding to 657 targets (including single proteins, protein families, protein complexes, organisms, cell lines and protein-protein interactions) and 18,319 compounds. For 145 of these targets, this represents the only source of information of bioactivity data in ChEMBL.

We examined the distribution of patent families among the different groups shown in Figure 1, to identify which group or groups were more or less likely to contain useful information, as this might facilitate the task of identifying the most useful patents in future analyses. The group that had the highest percentage of positive patents was group 1: 48 positive patent families in the 291 families examined (16.4%), followed by group 3: 88 positive families in the 1,912 examined (4.6%). There was no overlap between these 2 groups. Group 2 had 93 positive patent families, but 45 of them were already present in group 1. Group 4 had 95 positive patent families, but 86 of them were already present in group 3. There were very few patents with data in group 5 or group 6 (0.2 % and 0.5 % patent families of the examined ones, respectively) (table 1).

**Table 1.**

|  | Group 1 | Group 2 | Group 3 | Group 4 | Group 5 | Group 6 |
| --- | --- | --- | --- | --- | --- | --- |
| Total | 307 | 5,085 | 2,147 | 31,757 | 1,841,691 | 26,920 |
| Read | 291 | 3,655 | 1,912 | 2,736 | 1007 | 1017 |
| Positive | 48 | 93 | 88 | 95 | 2 | 5 |
| Negative | 243 | 3,562 | 1,824 | 2,641 | 1005 | 1012 |

As BindingDB also extracts data from patents, we were interested to examine the overlap between the two data sets. For all the targets found, we performed a search by target in BindingDB with the goal of comparing the results from the two different databases. Because BindingDB extracts only US granted patents, we used the patent family in order to do the comparison. We found 33 targets in both BindingDB and our method. Of the 70 patent families found by our method for these targets, 20 were also found in BindingDB. 50 families were found exclusively with our method, and 34 families were found exclusively by BindingDB.

### Examples of understudied targets with bioactivities found in patents

In the next section we briefly describe three specific examples of targets where we were able to identify and curate bioactivity data from the patent workflow described above.

### LATS1

LATS kinases are critical for organism fitness, genome integrity and cancer prevention. The core kinase module is evolutionarily conserved from yeast through flies to humans, although effectors and biological impact have expanded over the course of evolution^26^.

The family includes 2 members, LATS1 and LATS2, which play important roles in the control of tumor development and cell cycles through multiple mechanisms and signaling pathways, including those of p53, Hippo, and Wnt^27^. LATS1 phosphorylates Yes-associated protein (YAP) 1 and inhibits its translocation into the nucleus to regulate genes important for cell proliferation, cell death, and cell migration. Several recent studies have clearly established a role for the Hippo pathway in regulating cell contact inhibition, organ size control, and cancer development in mammals.

As a result of the current work 289 new molecules associated with this target were identified, 184 of which satisfy the IDG cut-off criteria. The source of these compounds is the patent US-20180344738-A1^28^, which describes molecules designed to promote cell proliferation, with applications such as chronic wound healing, promoting liver regrowth, or treating limbal stem cell deficiency. Of the molecules currently in ChEMBL associated with this target (extracted from 44 different papers), the most potent compounds are overwhelmingly from this one patent, albeit they do all contain the same core template.

### Histone-lysine N-methyltransferase SUV39H2

Suppressor of variegation 3-9 homolog 2 (SUV39H2, also known as KMT1B) is a member of the SUV39 subfamily of lysine methyltransferases (KMTs). It is an embryonic-specific protein and restricted to the testis in adult tissues of healthy individuals, localised in the nucleus^29^. SUV39H2 is primarily responsible for di- and tri-methylation of histone 3 lysine 9 (H3K9), which result in localized transcriptional silencing^30^. Deregulated H3K9me2/3 results in widespread abnormalities in various cellular functions such as cell proliferation, hypoxia, inflammation, cellular senescence, autophagy, and apoptosis. H3K9 methyltransferases have been implicated in several human diseases, including cancers, cocaine addiction, and HIV-1 latency. There is evidence that dysregulation of SUV39H2 contributes to carcinogenesis and is involved in invasion and metastasis of malignancy, and SUV39H2 is ubiquitously overexpressed in cancer tissues, such as leukemia, lymphomas, lung cancer, breast cancer, colorectal cancer, gastric cancer, and hepatocellular cancer. Some small molecule inhibitors against H3K9 methyltransferases have been developed, but only against some of them (SUV39H1, G9a, and GLP.39). No H3K9 methyltransferase inhibitors are currently in clinical trials^31^.

At the start of this work there were 19 molecules in ChEMBL with bioactivity data against SUV39H2, all of them from papers, but none of these molecules were within the IDG cutoff. Our patent workflow identified 460 molecules from just a single patent (US-20180273529-A1^32^), all within the corresponding IDG cutoff.

### G protein-coupled receptor 6

GPR6 is a constitutively highly active G-protein coupled receptor^33,34^. This receptor is mainly expressed in mammalian brain, specifically striatum and hypothalamus, with very low expression at peripheral level. Some endogenous ligands have been proposed for this receptor but due to inconsistencies among assays, it is categorized as orphan by the International Union of Basic and Clinical Pharmacology (IUPHAR)^35^. The physiological role of GPR6 has not been fully elucidated but it has been suggested that it could be involved in several diseases, such as Alzheimer’s disease, Parkinson’s disease, Huntington’s disease^36^ and schizophrenia.

187 molecules (from patents WO-2018183145-A1^37^ and EP-2882722-A1^38^) against GPR6 were identified from this work. These were complemented by an additional 130 molecules from BindingDB (from patent US-9181249-B2^39^). Figure 2 shows a timeline with patent and publication numbers by year for GPR6, showing that in this particular case significantly more data were reported via patent disclosures than in the scientific literature. Interestingly, a new clinical candidate (currently in phase 2 clinical trials) for Parkinson’s disease, CVN424 (a GPR6 inverse agonist), has been disclosed^40^. This molecule can be found in patents as early as 2015 (US-9181249-B2: “Substituted pyrido[3,4-b]pyrazines as GPR6 modulators”) but with reported bioactivities above the IDG criteria.

**Figure 2.**
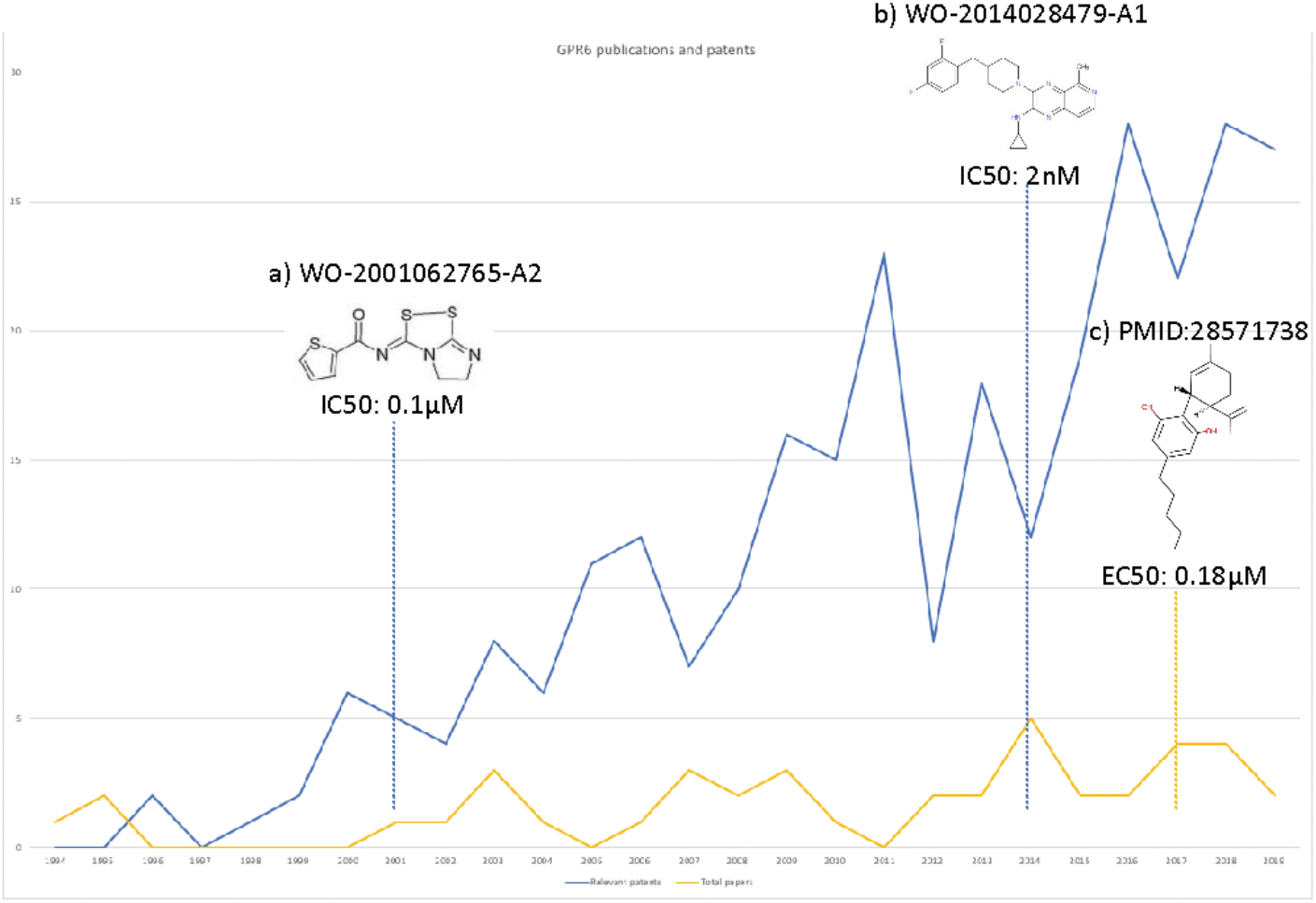
Numbers of relevant patents (blue) and publications (yellow) for GPR6 by year. Key compound disclosures in patents and literature indicated by dashed lines: a) Example of a compound reported in one of the earliest patents with data against GPR6. b) Example of a compound in one of the earliest patents identified by our method. c) First small molecule modulator reported in literature

## Discussion

The overall goal of this work was to identify bioactivity data on understudied targets from the patent literature, which could allow us to promote targets to the IDG Tchem category. We focused on small molecules only, but our workflow also identified several patents concerning antibodies or RNA as therapeutic agents. For the purposes of this work, we did not progress these patents further, but clearly they could also be useful in the context of “illuminating” new targets.

It should be noted that the work described here was conducted over a period of time, during which complementary data from other sources was added to the various resources concerned. This reflects the natural evolution of the underlying databases, each with its own update mechanism and release schedule procedures. Thus, for example, targets designated as Tdark or Tbio at the time when the research was initiated may have been promoted to a higher category based on separate evidence whilst the patent bioactivity work described here was underway. In the narrative below, the data reflects the particular snapshot corresponding to the time on which the work was initiated or completed, as appropriate.

Two of the proteins included in the list of targets that we searched for (sclerostin and exportin-1) were originally classified as Tbio at the start of this work, but later promoted to Tclin after the approval of the first in class drugs romosozumab and selinexor respectively.

In addition to these 2 targets that are now Tclin, 74 Tdark/Tbio targets were promoted to the Tchem category on the basis of the bioactivity data identified from our patent analysis. Coincidentally, over the same period of time, bioactivity data for 30 Tbio/Tdark targets was added into ChEMBL from the scientific literature. There was an overlap of 21 targets between the 2 sets. This shows the value of using patents as an additional source of bioactivity data.

The work described here involved manually reviewing many patents from the six groups described in Figure 1. As expected, the group with higher percentage of positive patents was Group 1 followed by Group 3. Groups 2 and 4 had lower percentages but still delivered useful and relevant patents. Clearly, for future work, it would be advantageous to develop methods that can reduce the level of manual review that is required. This is the focus on current ongoing work to develop machine learning models able to predict which patents should be prioritised for human examination and potential curation.

## Conclusions

Using relatively simple annotation and filtering pipelines, we have been able to identify a substantial number of patents containing quantitative bioactivity data for understudied targets that had not previously been reported in the peer-reviewed medicinal chemistry literature. This underlines the potential value in searching the patent corpus in addition to the more traditional peer-reviewed literature. The small molecules found in these patents, together with their measured activity against the targets are now accessible via the ChEMBL database and Pharos, and have contributed to the “illumination” of previously dark targets.

## Acknowledgements

This work was supported by US National Institutes of Health (NIH) grants U54 CA189205 and U24 224370 (Illuminating the Druggable Genome Knowledge Management Center (IDG KMC)) at the University of New Mexico, Novo Nordisk Foundation Center for Protein Research, European Bioinformatics Institute (EBI) and University of Miami.

The work was also in part supported by the Wellcome Trust [Grant numbers 104104/A/14/Z and 218244/Z/19/Z]. For the purpose of open access, the author has applied a CC BY public copyright licence to any Author Accepted Manuscript version arising from this submission. We also acknowledge funding from the Member States of the European Molecular Biology Laboratory.

We would like to thank Dr Barbara Zdrazil for helpful comments on this manuscript.

## References

1 Sheils TK, Mathias SL, Kelleher KJ, Siramshetty VB, Nguyen DT, Bologa CG et al. TCRD and Pharos 2021: mining the human proteome for disease biology. Nucleic Acids Res. 2021; 49:D1334–D1346.

2 Workman P, Collins I. Probing the Probes: Fitness Factors For Small Molecule Tools. Chem Biol 2010; 17: 561–577.

3 Garbaccio RM, Parmee ER. The Impact of Chemical Probes in Drug Discovery: A Pharmaceutical Industry Perspective. Cell Chem Biol 2016; 23: 10–17.

4 Mendez D, Gaulton A, Bento AP, Chambers J, De Veij M, Félix E et al. ChEMBL: towards direct deposition of bioassay data. Nucleic Acids Res. 2019; 47(D1):D930–D940.

5 Kim S, Chen J, Cheng T, Gindulyte A, He J, He S. et al. PubChem in 2021: new data content and improved web interfaces. Nucleic Acids Res 2021; 49: D1388–D1395

6 Gilson MK, Liu T, Baitaluk M, Nicola G, Hwang L, Chong J. BindingDB in 2015: A public database for medicinal chemistry, computational chemistry and systems pharmacology. Nucleic Acids Res 2016; 44: D1045–D1053

7 Southan, C, Várkonyi, P, Muresan, S. Quantitative assessment of the expanding complementarity between public and commercial databases of bioactive compounds. J Cheminform 2009; DOI: 10.1186/1758-2946-1-10 (https://jcheminf.biomedcentral.com/)

8 Senger, S. Assessment of the significance of patent-derived information for the early identification of compound–target interaction hypotheses. J Cheminform 2017; DOI: 10.1186/s13321-017-0214-2 (https://jcheminf.biomedcentral.com/)

9 Ashenden SK, Kogej T, Engkvist O, Bender A. Innovation in Small-Molecule-Druggable Chemical Space: Where are the Initial Modulators of New Targets Published? J Chem Inf Model 2017; 57:2741–2753

10 Akhondi SA, Klenner AG, Tyrchan C, Manchala AK, Boppana K, Lowe D et al. Annotated Chemical Patent Corpus: A Gold Standard for Text Mining. PLoS One 2014; DOI: 10.1371/journal.pone.0107477 (https://journals.plos.org/plosone/)

11 Suriyawongkul, I, Southan, C, Muresan, S. The Cinderella of Biological Data Integration: Addressing Some of the Challenges of Entity and Relationship Mining from Patent Sources. Conference: Data Integration in the Life Sciences, 7th International Conference, DILS 2010, Gothenburg, Sweden, August 25-27, 2010. Proceedings. 2010; DOI: 10.1007/978-3-642-15120-0\_9

12 Tyrchan C, Boström J, Giordanetto F, Jon Winter J, Muresan S. Exploiting Structural Information in Patent Specifications for Key Compound Prediction. J Chem Inf Model 2012; 52: 1480–1489

13 Fechete R, Heinzel A, Perco P, Mönks K, Söllner J, Stelzer G et al. Mapping of molecular pathways, biomarkers and drug targets for diabetic nephropathy. Proteomics Clin Appl 2011; DOI: 10.1002/prca.201000136 (https://onlinelibrary.wiley.com/journal/18628354)

14 Papadatos G, Davies M, Dedman N, Chambers J, Gaulton A, Siddle J et al. SureChEMBL: a large-scale, chemically annotated patent document database. Nucleic Acids Res 2016; 44: D1220–D1228

15 Gigani Y, Gupta S, Lynn A, Asotra K. Identification of BKCa channel openers by molecular field alignment and patent data-driven analysis. Pharm Biomed Res 2016; 2:22–29

16 Rodriguez-Esteban R, Bundschus M. Text mining patents for biomedical knowledge. Drug Discov Today 2016; 21: 997–1002

17 Kiss M, Nagy Á, Vincze, V, Almási, A, Alexin, Z, Csirik, J. A Manually Annotated Corpus of Pharmaceutical Patents. Text, Speech and Dialogue. TSD 2012. Lecture Notes in Computer Science. 2012; DOI: 10.1007/978-3-642-32790-2_16

18 Oprea TI, Bologa CG, Brunak S, Campbell A, Gan GN, Gaulton A et al. Unexplored therapeutic opportunities in the human genome. Nat Rev Drug Discov 2018; 17: 317–332

19 Amberger J, Bocchini CA, Scott AF, Hamosh A. McKusick’s Online Mendelian Inheritance in Man (OMIM). Nucleic Acids Res 2009; 37:D793–D796

20 Edwards AM, Isserlin R, Bader GD, Frye SV, Willson TM, Yu FH. Too many roads not taken. Nature 2011; 470: 163–165

21 Armstrong JF, Faccenda E, Harding SD, Pawson AJ, Southan C, Sharman JL et al. The IUPHAR/BPS Guide to PHARMACOLOGY in 2020: extending immunopharmacology content and introducing the IUPHAR/MMV Guide to MALARIA PHARMACOLOGY. Nucleic Acids Res 2020; 48:D1006–D1021

22 Avram S, Bologa CG, Holmes J, Bocci G, Wilson TB, Nguyen DT et al. DrugCentral 2021 supports drug discovery and repositioning. Nucleic Acids Res 2021; 49:D1160–D1169

23 https://www.wipo.int/classifications/ipc/en/

24 https://worldwide.espacenet.com/classification

25 Gaulton A, Kale N, van Westen GJP, Bellis LJ, Bento AP, Davies M et al. A large-scale crop protection bioassay data set. Sci Data 2015; DOI:10.1038/sdata.2015.32 (https://www.nature.com/sdata/)

26 Furth N, Aylon Y. The LATS1 and LATS2 tumor suppressors: beyond the Hippo pathway. Cell Death Differ 2017; 24:1488–1501

27 Xu B, Sun D, Wang Z, Weng H, Wu D, Zhang X et al. Expression of LATS family proteins in ovarian tumors and its significance. Hum Pathol 2015; 46: 858–867

28 6-6 Fused Bicyclic Heteroaryl Compounds and their Use as LATS Inhibitors. Behnke D et al. Novartis AG. US-20180344738-A1.

29 Li B, Zheng Y, Yang L. The Oncogenic Potential of SUV39H2: A Comprehensive and Perspective View. J Cancer. 2019; 10:721–729

30 Kaniskan HÜ, Martini ML, Jin J. Inhibitors of Protein Methyltransferases and Demethylases. Chem Rev 2018; 118: 989–1068

31 Saha N, Muntean AG. Insight into the multi-faceted role of the SUV family of H3K9 methyltransferases in carcinogenesis and cancer progression. Biochim Biophys Acta Rev Cancer 2021; DOI:10.1016/j.bbcan.2020.188498 (https://www.sciencedirect.com/journal/biochimica-et-biophysica-acta-bba-reviews-on-cancer)

32 Bicyclic compound and use thereof for inhibiting SUV39H2. Matsuo Y et al. Oncotherapy Science Inc. US-20180273529-A1.

33 Morales P, Isawi I, Reggio PH. Towards a better understanding of the cannabinoid-related orphan receptors GPR3, GPR6, and GPR12. Drug Metab Rev. 2018; 50:74–93.

34 Sun H, Monenschein H, Schiffer HH, Reichard HA, Kikuchi S, Hopkins M et al. First-Time Disclosure of CVN424, a Potent and Selective GPR6 Inverse Agonist for the Treatment of Parkinson’s Disease: Discovery, Pharmacological Validation, and Identification of a Clinical Candidate. J Med Chem 2021; 64: 9875–9890

35 Alexander SP, Battey J, Benson HE, Benya RV, Bonner TI, Davenport AP et al. Class A Orphans (version 2019.5) in the IUPHAR/BPS Guide to Pharmacology Database. GtoPdb CITE 2019; DOI: 10.2218/gtopdb/F16/2019.5

36 Isawi IH, Morales P, Sotudeh N, Hurst DP, Lynch DL, Reggio PH. GPR6 Structural Insights: Homology Model Construction and Docking Studies. Molecules 2020; DOI: 10.3390/molecules25030725.

37 Piperidinyl- and piperazinyl-substituted heteroaromatic carboxamides as modulators of GPR6. Green J et al. Takeda Pharmaceuticals Company Limited. WO-2018183145-A1.

38 Quinoxaline derivatives as GPR6 modulators. Hitchcock S et al. Envoy Therapeutics Inc. EP-2882722-A1.

39 Substituted pyrido[3,4-b]pyrazines as GPR6 modulators. Brown J et al. Takeda Pharmaceutical Company Limited. US-9181249-B2.

40 Brice NL, Schiffer HH, Monenschein H, Mulligan VJ, Page K, Powell J et al. Development of CVN424: a novel GPR6 inverse agonist for Parkinson’s disease. J Pharmacol Exp Ther 2021; DOI: 10.1124/jpet.120.000438 (https://jpet.aspetjournals.org/)

